## Supplementary Information with embedded figures for "ADAP1 promotes latent HIV-1 reactivation by selectively tuning a T cell signaling-transcriptional axis"

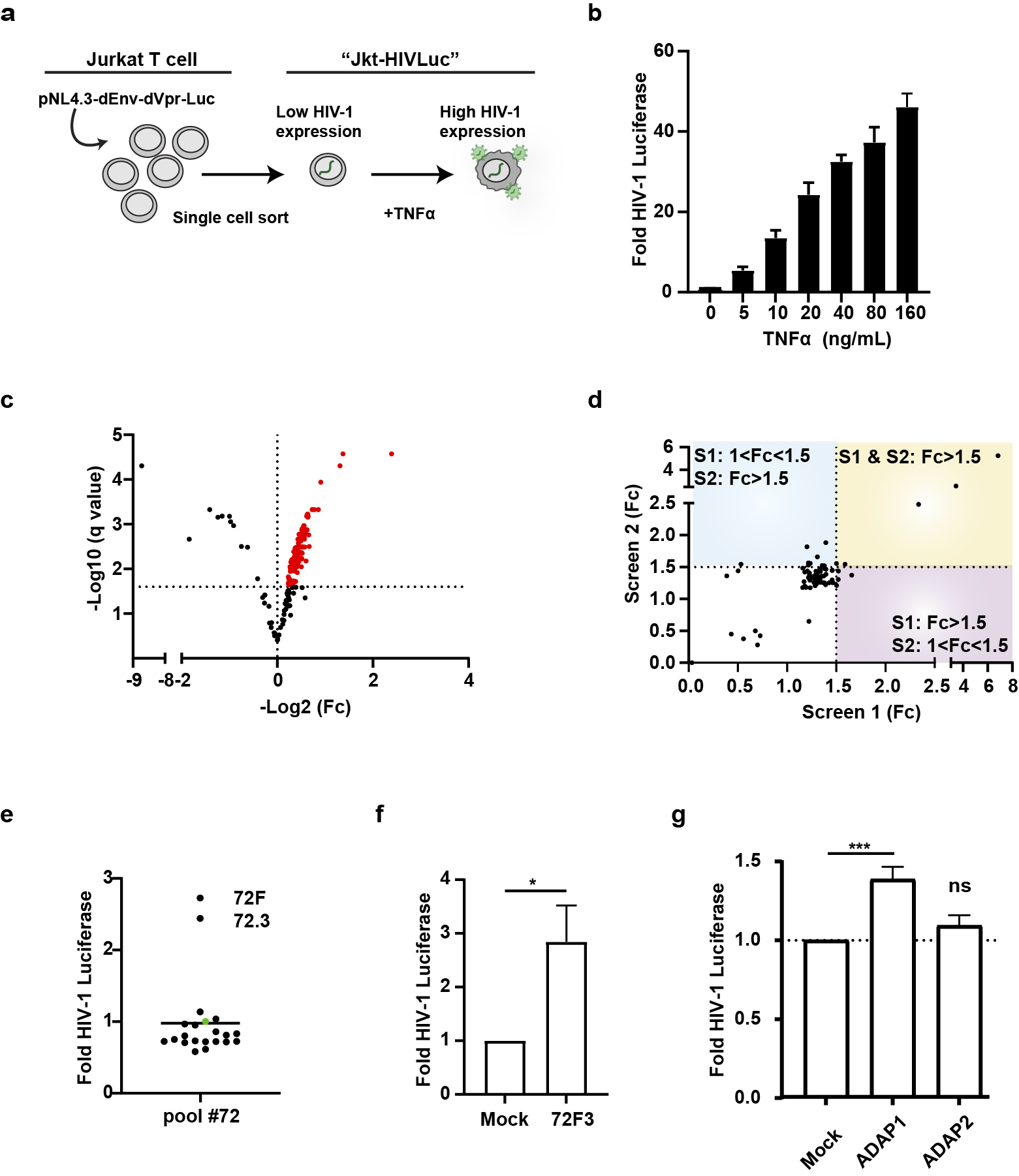

**Extended Data Figure 1. Development of Jkt-HIVLuc, a cell-based model of latency used in the gain-of-function screen to identify undescribed latent HIV-1 activating factors.**

a) Schematic of the development of Jkt-HIVLuc cell-based model. Jurkat T cells were transduced with pNL4.3-delta*Env*- delta*Vpr*-Luc-VSVG. Cells were single cell sorted and clonally expanded. Clones were screened by TNFα treatment (25 ng/mL for 16 hrs) to identify those with latent-reactivation switching capabilities.
b) Selected Jkt-HIVLuc clone responds to TNFα in a dose-dependent manner exhibiting increasing luciferase activity. Mean ± S.D. fold luciferase activity compared to no treatment (n=3).
c) Volcano plot of second screen to validate first screen (**Fig. 1d**) using different batch of generated lentiviral pools. Results are a summary of pooled samples and statistical significance (determined by multiple unpaired t test) for phase 1 gain-of-function screen. Each dot is a pool sample (representing 96 cDNAs) mean luciferase activity (n=3). Dots in red were above the chosen cutoff of Log_2_Fc > 0 with an FDR < 2.5%.
d) Selection criteria for determining which pools to prioritize for decomposition. Pools with an FDR < 2.5% and Log_2_Fc > 0 from both screens (**Fig. 1d, Extended Data Fig. 1c**) were compared. Pools selected included: (the yellow quadrant) pools that induced an HIV-1 Luciferase Fc > 1.5 in both screens; (the pink quadrant) pools that induced an HIV-1 Luciferase Fc > 1.5 in screen 1 but still had an Fc > 1 in screen 2; (the blue quadrant) pools that induced an HIV-1 Luciferase Fc > 1.5 in screen 2 but still had an Fc > 1 in screen 2.
e) Representative example of screen phase 2 using pool number 72 to identify activating factor within the pool. Each dot represents the mean ± S.D. fold luciferase activity (n=4) of phase 2 pools (A-H rows= 12 cDNA each, or 1-12 columns= 8 cDNAs each). The mean activity is represented by the line and the green dot denotes mock control.
f) Representative example of screen phase 3 validating the intersecting well of positive pools from phase 2 (**Extended Data Fig. 1e**) contain activating factors. Data represent mean ± S.D. fold of 4 independent experiments (n = 3) normalized to mock. [Paired t-test comparing mock and sample independent experiments]. *P < 0.05, **P < 0.01, ***P < 0.001, ns= not significant.
g) Fold luciferase activity of Jkt-HIVLuc cells transduced with pTRIP lentiviruses expressing ADAP1 or ADAP2. Data represents mean ± S.D fold luciferase activity (n = 3). [one-way ANOVA with comparison to mock]. *P< 0.05, **P< 0.01, ***P< 0.001, ns= not significant.

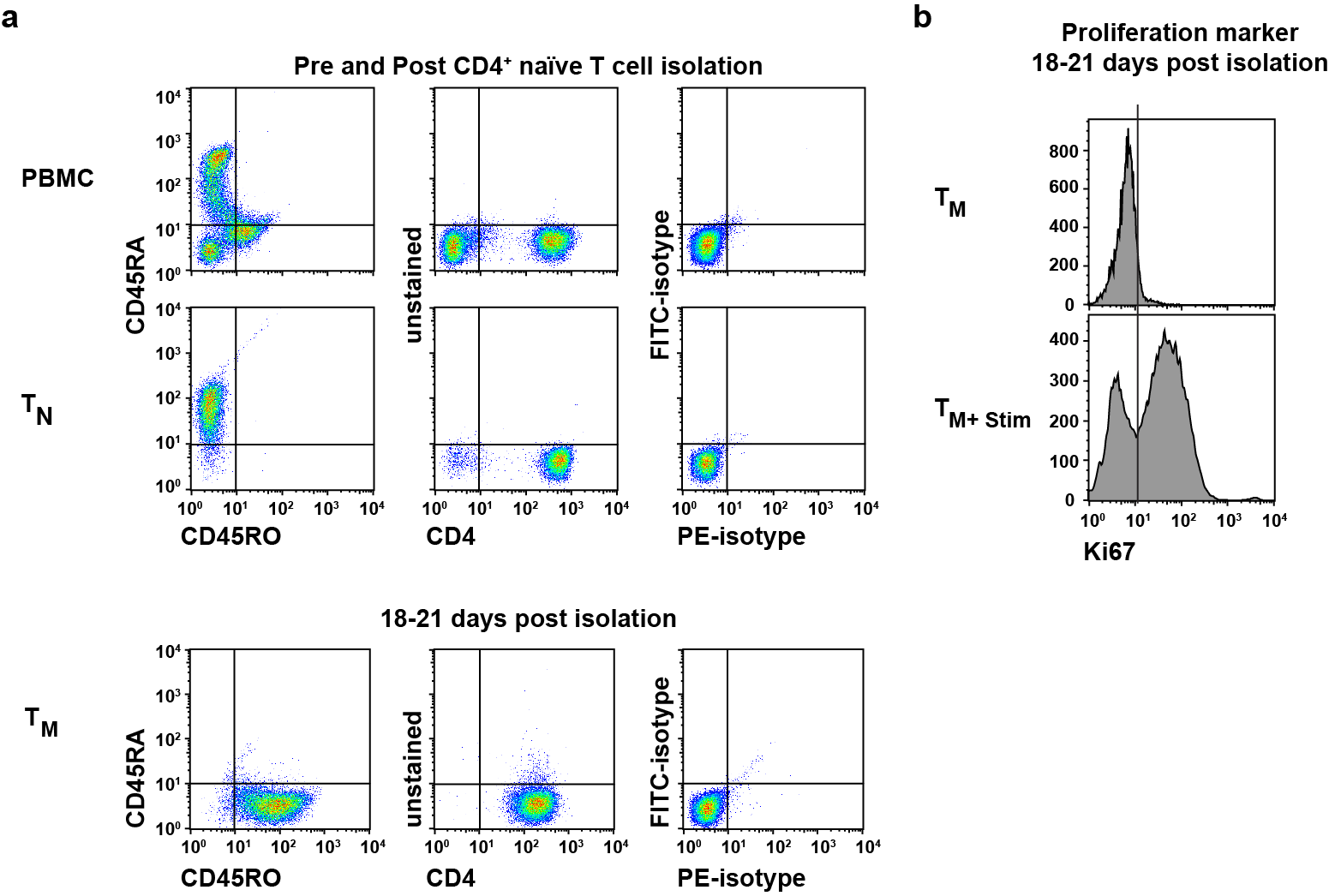

**Extended Data Figure 2. Analysis of primary T cell states.**

a) Representative flow cytometry analysis comparing purity after naïve CD4^+^ T cell isolation from PBMC yielding a CD4^+^CD45RA^+^CD45RO^-^ population. Approximately 18-21 days post isolation and effector-to-memory transition, population yielded a CD4^+^CD45RA^-^CD45RO^+^ phenotype.
b) Representative flow cytometric analysis measuring intracellular proliferation marker Ki67 comparing cells that have fully transitioned into resting, non-proliferating memory T cells (T_M_), and stimulated cells (T_M_+Stim).

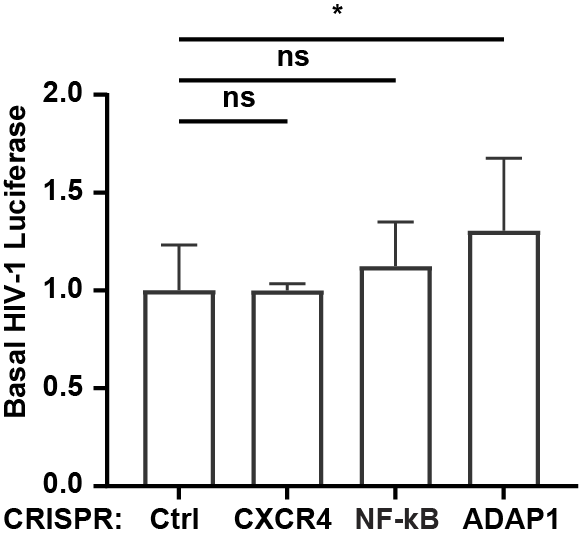

**Extended Data Figure 3. Primary resting ADAP1^CRISPR^ T cells have increased basal HIV-1 activity.**

Luciferase analysis of resting HIV-Ctrl^CRISPR^, HIV-CXCR4^CRISPR^, HIV-NFκB^CRISPR^, and HIV-ADAP1^CRISPR^ T_M_ in the absence of stimulation. For all samples, data represents mean ± S.D. raw luciferase units of 3 donors (n = 3 each) normalized to HIV-Ctrl^CRISPR^ (Ctrl = 1). [one-way ANOVA test with multiple comparison to Ctrl^CRISPR^]. *P< 0.05, **P< 0.01, ***P< 0.001, ns= not significant.

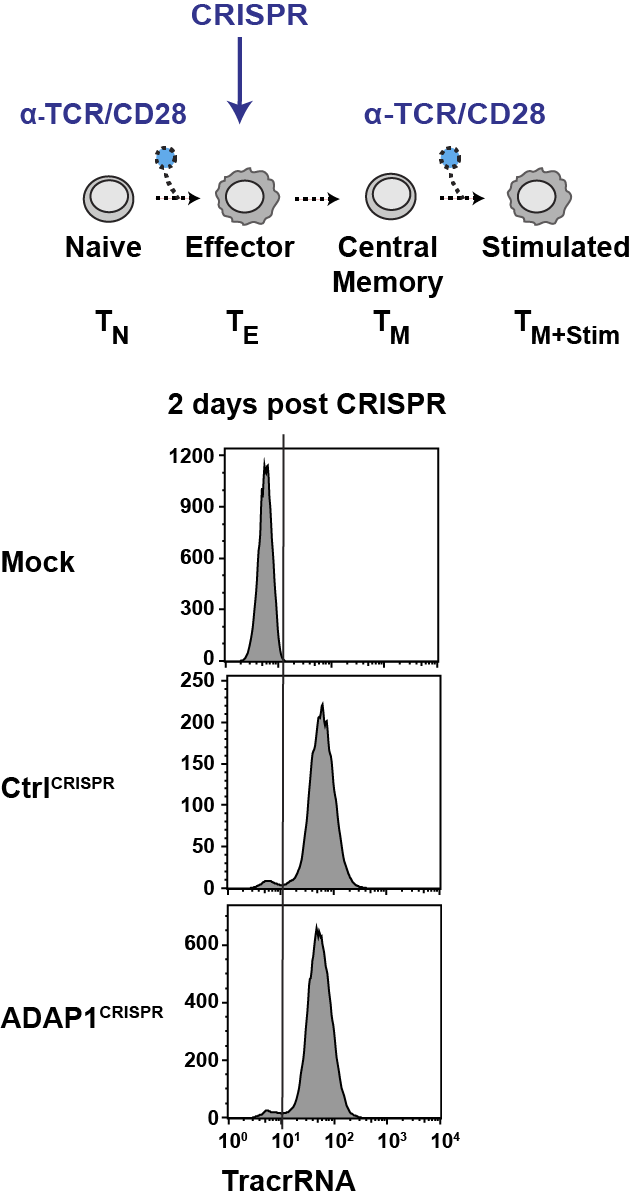

**Extended Data Figure 4. Validation of CRISPR-Cas9 ribonucleoprotein electroporation in generation of ADAP1^CRISPR^ and Ctrl^CRISPR^ cells.**

Schematic of CRISPR-Cas9 schedule for generating ADAP1^CRISPR^ and Ctrl^CRISPR^ T_M_. A ribonucleoprotein (RNP) complex consisting of Cas9 complexed with a fluorescent tracrRNA (ATTO550) and gRNA were assembled before delivering to cells via electroporation. Two days post electroporation, the fluorescent tracrRNA ATTO550 was used to monitor delivery efficiency of the RNP by flow cytometry.

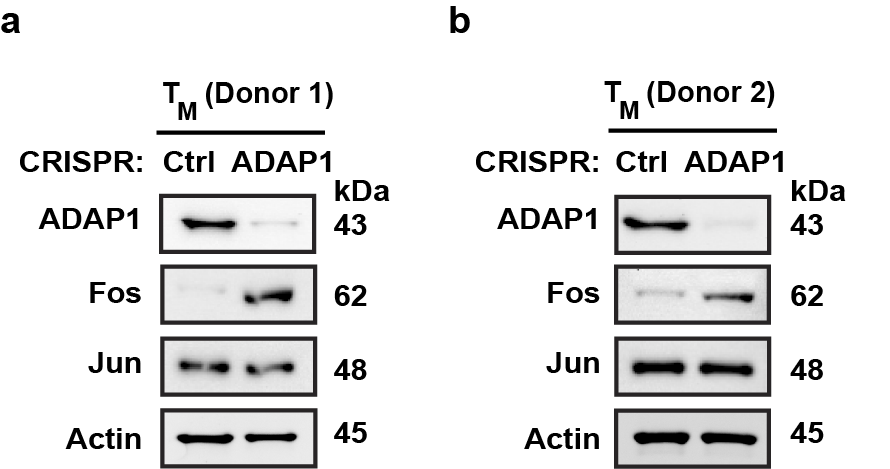

**Extended Data Figure 5. Primary resting ADAP1^CRISPR^ T cells have elevated basal Fos expression.**

a) and **b)** Representative western blot analysis of resting Ctrl^CRISPR^ and ADAP1^CRISPR^ T_M_ in unstimulated donors.

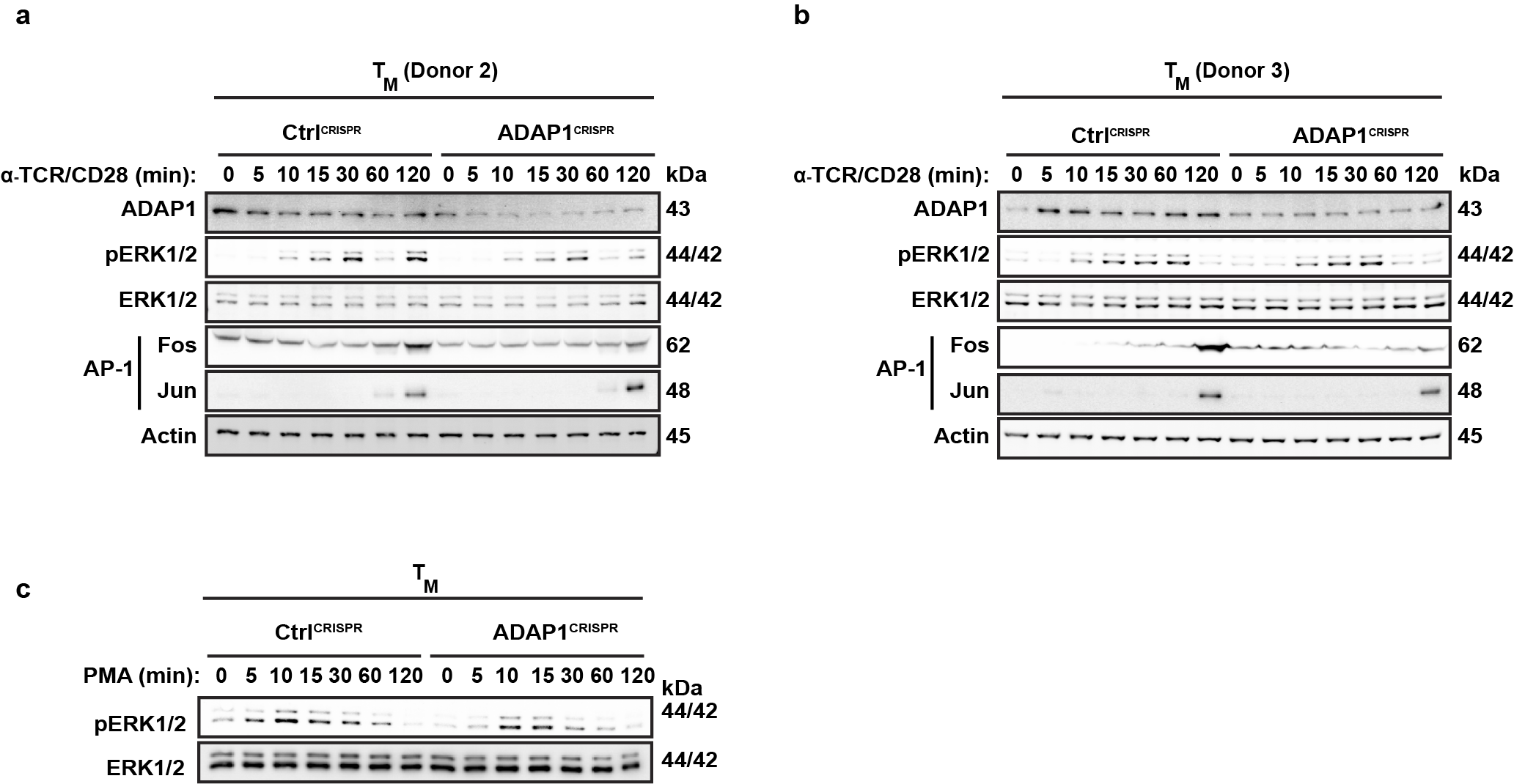

**Extended Data Figure 6. Loss of *ADAP1* in primary human cells impairs ERK–AP-1 activation.**

a) and **b)** Additional donors to complement **Fig. 6f**. Western blot analysis of primary ADAP1^CRISPR^ and Ctrl^CRISPR^ T_M_ stimulated with anti-TCR/anti-CD28 beads in a time course-dependent manner from 0 to 120 min.
c) Western blot analysis of primary ADAP1^CRISPR^ and Ctrl^CRISPR^ T_M_ stimulated with 25 ng/mL PMA in a time course-dependent manner from 0 to 120 min.

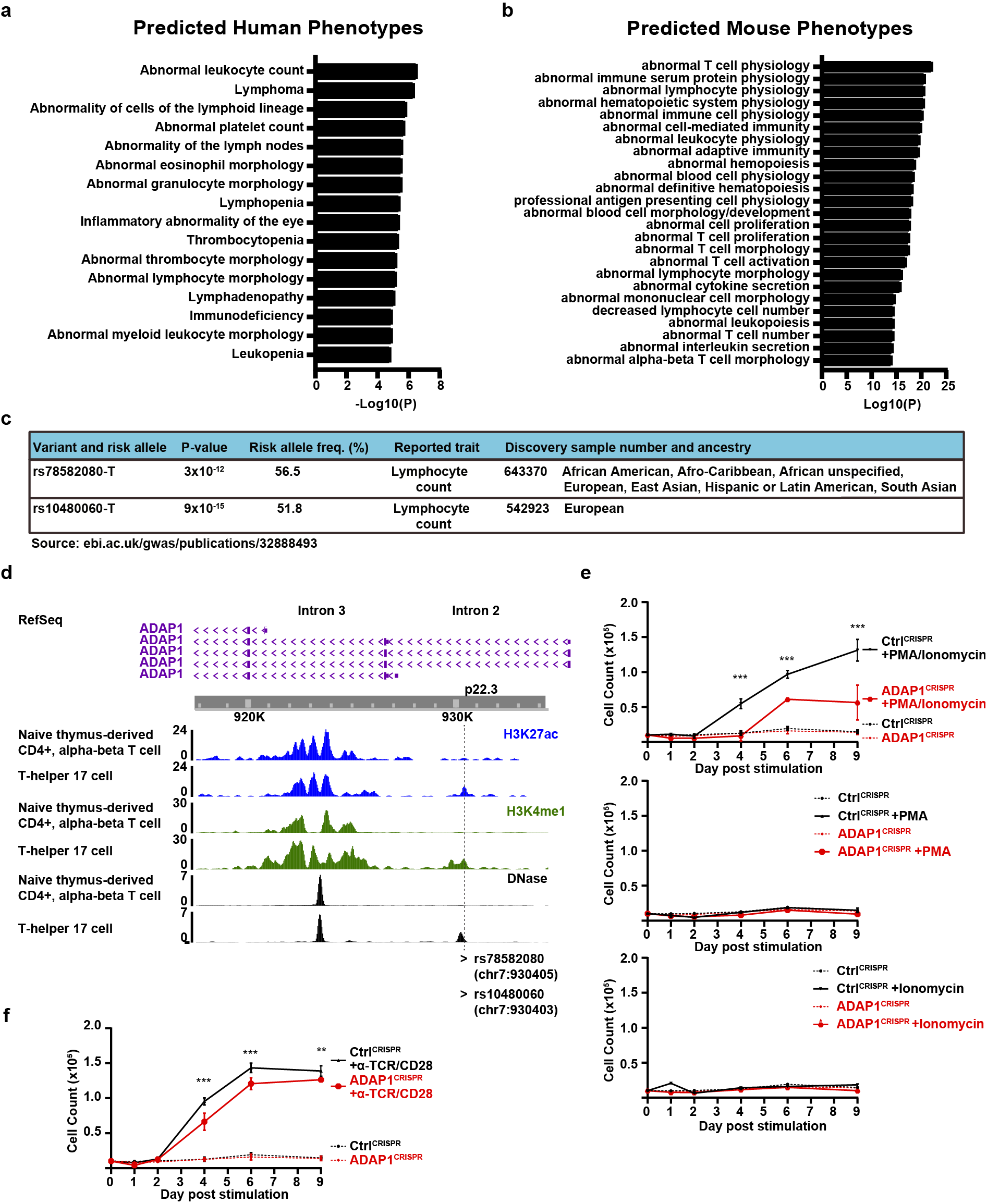

**Extended Data Figure** **7. Loss of *ADAP1* results in reduced T cell proliferation *ex vivo* consistent with GWAS linking non-coding SNPs in putative enhancers to altered T lymphocyte count trait.**

a) Predicted human phenotypes and **b)** mouse phenotypes based on induced genes found in both ADAP1^CRISPR^ and Ctrl^CRISPR^ samples sets. Analysis performed using ToppGene Suites^52^.
c) *ADAP1* SNPs information sourced from publicly available GWAS <https://www.ebi.ac.uk/gwas/studies/GCST90002320>^20^.
d) ChIP-seq and DNase-seq tracks sourced from publicly available datasets using WashU Epigenome Browser <https://epigenomegateway.wustl.edu/>^53^. Tracks reflect H3K27ac (blue), H3K4me1 (green), and DNase (black) of naïve CD4^+^ T cells and Th17 effector cells. SNPs ID and locations are labeled on the bottom and depicted with vertical dashed line. Reference sequences for main transcripts are depicted above in purple along with intron reference.
e) Cell counting analysis of Ctrl^CRISPR^ and ADAP1^CRISPR^ T_M_ stimulated with 25 ng/mL PMA and/or 1 μM ionomycin in the absence of exogenous growth factor IL-2. Data represents mean ± S.D. number of cells (n=3). [two-way ANOVA test with multiple comparison]. *P < 0.05, **P < 0.01, ***P < 0.001, ns= not significant
f) Cell counting analysis of Ctrl^CRISPR^ and ADAP1^CRISPR^ T_M_ stimulated with anti-TCR/anti-CD28 beads in the absence of exogenous growth factor IL-2. Data represents mean ± S.D. number of cells (n=3). [two-way ANOVA test with multiple comparison]. *P < 0.05, **P < 0.01, ***P < 0.001, ns= not significant.

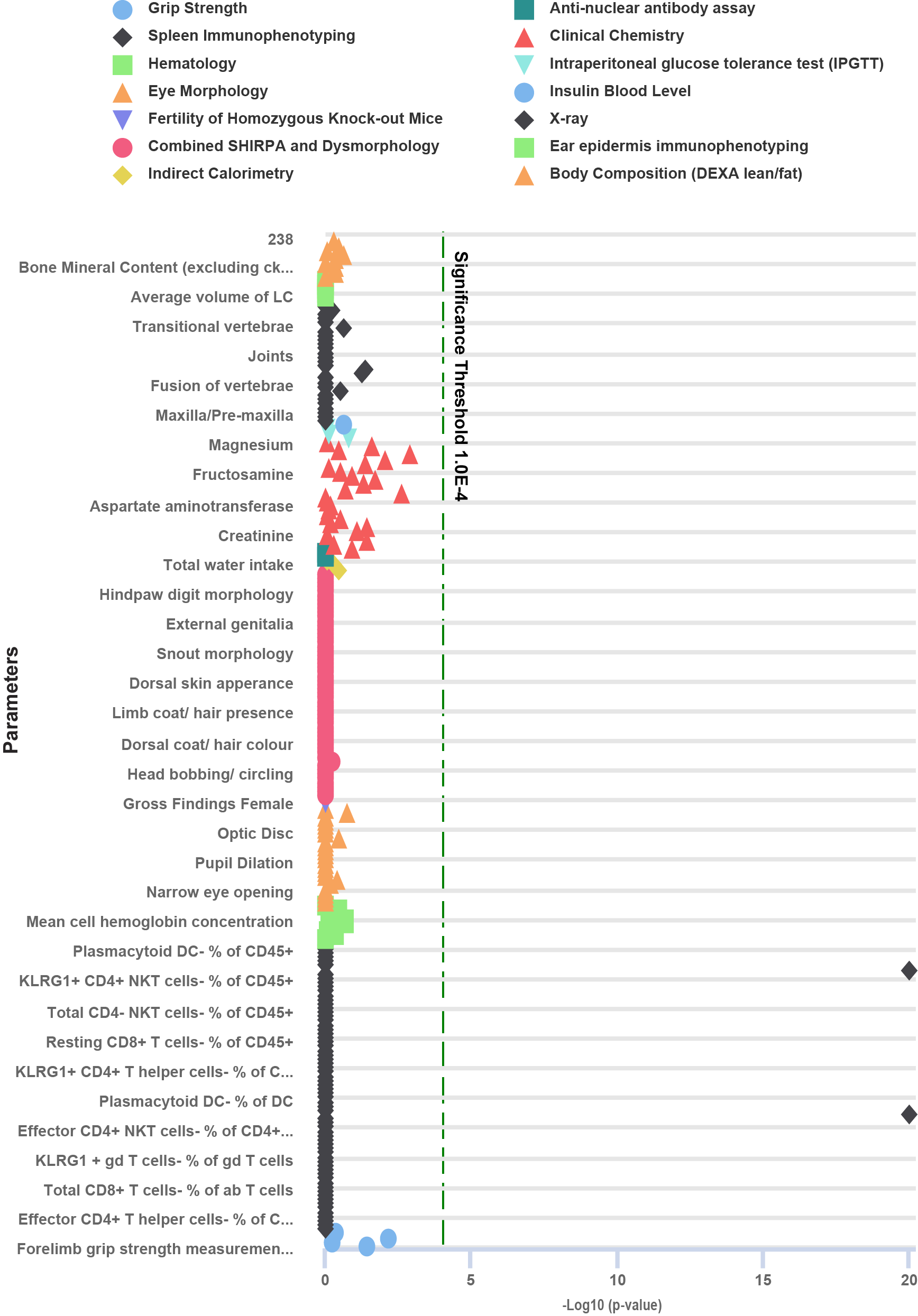

**Extended Data Figure** **8. *ADAP1* mutant mice from International Mouse Consortium Project have impaired NK cell counts.**

Phenotypic analysis on unchallenged *ADAP1* mutant mouse line (Adap1^tm1a(EUCOMM)Wtsi^). Sourced from International Mouse Consortium Project [www.mousephenotype.org](http://www.mousephenotype.org)^43^.
**Supplementary Table 1. ADAP1 IP and mass spectrometry dataset corresponding to donor in Fig. 4b.** File added separately (.xlsx)

**Supplementary Table 2. ADAP1 IP and mass spectrometry dataset corresponding to donor in Fig. 4c-d.** File added separately (.xlsx)

**Supplementary Table 3. Average Log_2_Fc of DEG of stimulated vs unstimulated primary T cells corresponding to scRNAseq in Fig. 5c-h.** File added separately (.xlsx)
